# Chicken peripheral blood lymphocyte response to ALV-J infection assessed by single-cell RNA sequencing

**DOI:** 10.1101/2021.01.12.426350

**Authors:** Manman Dai, Min Feng, Ziwei Li, Weisan Chen, Ming Liao

## Abstract

Chicken peripheral blood lymphocytes (PBLs) exhibit wide-ranging cell types, but current understanding of their subclasses, immune cell classification, and function is limited and incomplete. Previously, we found that viremia caused by avian leukosis virus subgroup J (ALV‐J) was eliminated by 21 days post infection (DPI), accompanied by increased CD8^+^ T cell ratio in PBLs and low antibody levels. Here we performed single-cell RNA sequencing (scRNA-seq) of PBLs in ALV-J infected and control chickens at 21 DPI to determine chicken PBL subsets and their specific molecular and cellular characteristics, before and after viral infection. Eight cell clusters and their potential marker genes were identified in chicken PBLs. T cell populations (clusters 6 and 7) had the strongest response to ALV-J infection at 21 DPI, based on detection of the largest number of differentially expressed genes (DEGs). T cell populations of clusters 6 and 7 could be further divided into four subsets: activated CD4^+^ T cells (cluster A0), Th1-like cells (cluster A2), Th2-like cells (cluster A1), and cytotoxic CD8^+^ T cells. Hallmark genes for each T cell subset response to viral infection were initially identified. Furthermore, pseudotime analysis results suggested that chicken CD4^+^ T cells could potentially differentiate into Th1-like and Th2-like cells. Moreover, ALV-J infection probably induced CD4^+^ T cell differentiation into Th1-like cells in which the most immune related DEGs were detected. With respect to the control group, ALV-J infection also had an obvious impact on PBL cell composition. B cells showed inconspicuous response and their numbers decreased in PBLs of the ALV-J infected chickens at 21 DPI. Percentages of cytotoxic Th1-like cells and CD8^+^ T cells were increased in the T cell population of PBLs from ALV-J infected chicken, which were potentially key mitigating factors against ALV-J infection. More importantly, our results provided a rich resource of gene expression profiles of chicken PBL subsets for a systems-level understanding of their function in homeostatic condition as well as in response to viral infection.

## INTRODUCTION

Adaptive immunity is known to play a vital protective role against avian viral infections. However, studies so far have focused on the avian innate immune response, while research on avian T cell or B cell immunity is still in its infancy [1]. For example, many important marker genes of the chicken immune cells are unknown, including effector or memory T cells and B cells; this greatly limit the immune cell phenotyping and subsequent immune function and mechanistic studies. Specifically, effector CD4^+^ T cells can differentiate into many T helper (Th) subsets, resulting in the production of different cytokine and effector functions. The Th1-Th2 paradigm is reported to exist in chickens [2]. However, whether this paradigm holds true at the cellular and molecular levels and whether chicken Th cells can become terminally polarized to a Th1 or Th2 phenotype remain to be verified. Furthermore, it is also important and interesting to thoroughly characterize the molecular signatures of the CD4^+^ T, CD8^+^ T cells and B cells during homeostasis and after pathogen exposure.

Chicken peripheral blood lymphocytes (PBLs), which contain various lymphocytes including T cells, B cells and other cell types, are reported to execute important functions in eliminating avian viral infections [3–5]. However, such studies are usually based on bulk PBLs measurements overlooking the complexity of diverse cell types. Recent advances in single-cell RNA-seq (scRNA-seq) allow the breakdown of complex tissues or host compartments into individual cell types for exploring their relevance in health and disease [6]. scRNA-seq has already been used to investigate the immune response of human peripheral blood cells under infectious of pathogenic microorganisms including salmonella, severe acute respiratory syndrome corona virus 2 (SARS-CoV-2), and influenza infection[7–9]. In chickens, the cell lineage characteristics in some tissues have been identified using scRNA-seq, such as the developing chicken limb [10], chicken skeletal muscle [11], and embryonic chicken gonad [12]. However, no reported study using scRNA-seq to determine chicken immune cell subsets or lineages. Moreover, to our knowledge, scRNA-seq technology has not even been applied to study chicken PBL responses to any viral infection.

Avian leukosis virus subgroup J (ALV‐J), an avian oncogenic retrovirus, causes enormous economic losses in the global poultry industry as there are currently no vaccines or drug treatments [13]. A potential vaccine for ALV-J has been reported to induce significantly increased CD4^+^ and CD8^+^ T cell percentage as well as IL-4 and IFN-γ levels in immunized chickens [14]. Unfortunately, there have been few studies exploring the specific T cell functions against ALV. A full understanding of the ALV-specific cellular immune response in chickens is likely the premise for developing effective vaccines. In our previous study, we found that ALV‐J viremia was eliminated by 21 days post infection (DPI) when a significantly up-regulated CD8^+^T cell ratio and a very low serum antibody level in the peripheral blood were detected [3]. In fact, PBLs contains many cell types besides CD8^+^ T cells and B cells. Usually, these immune cells are able to form a complex network of communications that maintains an orchestrated and dynamic immune response to eliminate invading pathogens [9]. Hence, elucidation of the response of different chicken PBL subsets to viral infection is worthwhile.

In the current study, we performed 10x scRNAseq on PBLs in ALV-J infected and uninfected chickens at 21 DPI to comprehensively identify PBL subsets and characterize their specific cellular and molecular responses after viral infection. More importantly, we provide evidence to verify that chicken Th cells can terminally polarized to a Th1 or Th2 phenotype. Moreover, we developed an extensive catalogue of candidate marker genes or immune response factors for functional follow-up studies to identify or complement known and unknown chicken immune cells and their function.

## MATERIALS AND METHODS

### Sample preparation

Samples from ALV-J infected #12 PBL and control #29 PBL at 21 DPI were prepared as previously described [3]. Briefly, four‐week‐old specific‐pathogen‐free (SPF) chickens were infected with ALV-J, and the virus was eliminated at 21 DPI when a significantly up-regulated CD8^+^ T cell ratio in PBLs was detected compared to the control group [3]. To further explore the immune response of various immune cells in PBLs, we further investigated the single-cell survey of the chicken PBL response to ALV-J infection at 21 DPI.

### Single cell suspension for 10x scRNAseq

PBLs of ALV-J infected #12 and control #29 chickens at 21 DPI were, respectively, resuspended in PBS (calcium and magnesium-free; Gibco, USA) with 0.4% bovine serum albumin (BSA; Solarbio, China), followed by filtering through a 40 μm cell strainer (Biosharp, China). Cell concentration and viability were assessed using Trypan Blue and a Neubauer hemocytometer (Sigma‐Aldrich, St. Louis, MO, USA). Cell viability in both samples was about 80%. Subsequently, the cell density was adjusted to 1 × 10^6^ cells/mL. High quality single cell suspension were subjected to encapsulation using a 10x Genomics v.3 kit (10x Genomics, USA).

### Library preparation for 10x scRNAseq

Single cell encapsulation, complementary DNA (cDNA) library synthesis, RNA-sequencing, and data analysis were completed by Gene Denovo (Guangzhou, China). The single-cell suspensions were bar-coded and reverse-transcribed into scRNA-seq libraries using the Chromium Single Cell3’ Gel Bead-in Emulsion (GEM) Library and Gel Bead Kit (10x Genomics) according to the manufacturer’s protocol. Briefly, single cells of ALV-J infected #12 PBL and control #29 PBL at 21 DPI were respectively barcode-labeled and mixed with reverse transcriptase into GEMs; the cDNA library was then using PCR with the sequencing primers R1 and R2, and subsequently ligated to Illumina sequencing adapters with P5 and P7. Finally, the cDNA libraries were sequenced on the Illumina 10x Genomics Chromium platform (Illumina Novaseq 6000). An average of 18 895 reads per cell in #12 PBL and 30 540 mean reads per cell in #29 PBL were obtained, respectively.

### ScRNA-seq data processing and analysis

#### (1) Data processing

Cell Ranger (http://support.10xgenomics.com/single-cel/software/overview/welcome) uses an aligner called STAR (https://github.com/alexdobin/STAR), which performs splicing-aware alignment of reads to the chicken ENSEMBL genome assembly and annotation with the latest genome version of *Gallus gallus* (GRCg6a) [15]. Only reads that are confidently mapped to the transcriptome are used for Unique Molecular Identifier (UMI) counting. Cells with unusually high numbers of UMIs (≥8000) or mitochondrial gene percent (≥10%) were filtered out. Cells with <500 or >4000 were also excluded. Using the R package Seurat v.2.3.2 [16], UMI counts were then Log-normalized and any variation due to the library size or mitochondrial UMI count percentage was then regressed via a variance correction using the function ScaleData.

#### (2) Dimensionality reduction and visualization

Significant principal components (PCs) were determined for each sample as those that had a strong enrichment of low p-value genes for downstream clustering and dimensional reduction following the jackStraw procedure [16]. Then, Seurat was used to implement the graph-based clustering approach [17,18]. T-distributed Stochastic Neighbor Embedding (t-SNE) [19] or Uniform Manifold Approximation and Projection (UMAP) [20] in Seurat were used to visualize and explore these datasets.

#### (3) Differentially expressed gene (DEG) (up-regulation) analysis per cluster

We used the Wilcoxon rank sum test [21] to identify differential expression for a single cluster, compared to all other cells. We identified differentially expressed genes (DEGs) according to the following criteria: 1) *P* value ≤ 0.01; 2) logFC ≥ 0.360674 (logFC means log fold-change of the average expression between the two compared groups); 3) The percentage of cells where the gene is detected in a specific cluster > 25%. Gene ontology (GO) enrichment analysis selects all GO terms that are significantly enriched in DEGs compared to the genome background; furthermore, the DEGs are filtered to correspond to biological functions. All DEGs were mapped to GO terms in the Gene Ontology database [22], gene numbers were calculated for every term, and significantly enriched GO terms in DEGs compared to the genome background were identified by hypergeometric testing. Finally, Pearson’s correlation analysis was performed to investigate correlations between different clusters based on the levels of gene expression.

#### (4) Marker gene analysis

We further selected the top five expressed genes as marker genes according to the result of differentially expressed genes. The expression distribution of each marker gene was then demonstrated using bubble diagrams.

### Pseudo temporal ordering of cells

Single cell trajectory was analyzed using a matrix of cells and gene expressions in Monocle 2 (v.2.6.4) [23]. Monocle reduces the space in which cells are embedded to two dimensions and orders the cells (parameters used: sigma = 0.001, lambda = NULL, param.gamma = 10, tol = 0.001). Once the cells were ordered, the trajectory (with a tree-like structure, including tips and branches) could be visualized in the reduced dimensional space.

### DEG analysis in cell clusters of PBLs from the ALV-J infected and control chickens at 21 DPI

To explore the response of each cluster in PBL, we further analyzed the DEGs in cell clusters of PBLs from the infected and control chickens using Seurat’s R package. A hurdle model in MAST (Model-based Analysis of Single-cell Transcriptomics) [24] was used to identify DEGs group in one cluster. DEGs between the ALV-J infected and control samples were identified by the following criteria: 1) |log2FC| ≥ 1; 2) *P* value ≤ 0.01; and 3) percentage of cells in which the gene was detected in a specific cluster > 25%. Identified DEGs were subsequently subjected to GO enrichment analysis as described above.

### Protein-protein interaction network analysis

The interaction network of the candidate DEGs was constructed using String v.10.0 and Cytoscape (v.3.3.0) software. Specifically, the protein-protein interaction network was identified using String [25], which determined genes as nodes and interactions as lines in a network. The final network file was visualized using Cytoscape software [26] to present a core and hub gene biological interaction.

## Results

### Single-cell transcriptomics identified eight distinct cell clusters in the PBLs collected from ALV-J-infected and control chickens at 21 DPI

We used the 10x Genomics platforms to perform 3’ scRNA-seq on PBLs collected from ALV-J infected and PBS-treated control chickens at 21 DPI, respectively. Details on the statistics of scRNA-seq are summarized in file S1. A total of 13 766 cells in the PBLs from ALV-J infected chicken and 9786 cells in the control PBLs were profiled, and eight distinct clusters were obtained and visualized using UMAP (Fig. 1). Clusters 6, 7, and 8 occupied a very small percentage in chicken PBLs (Fig. 1C), and the percentage of clusters 0, 1, and 3 were increased in the PBLs from ALV-J infected chicken when compared to the control. Conversely, the percentage of clusters 2, 6, 7, and 8 were distinctly reduced in the PBLs from ALV-J infected chicken at 21 DPI (Fig. 1 A-C).

**Figure 1.**
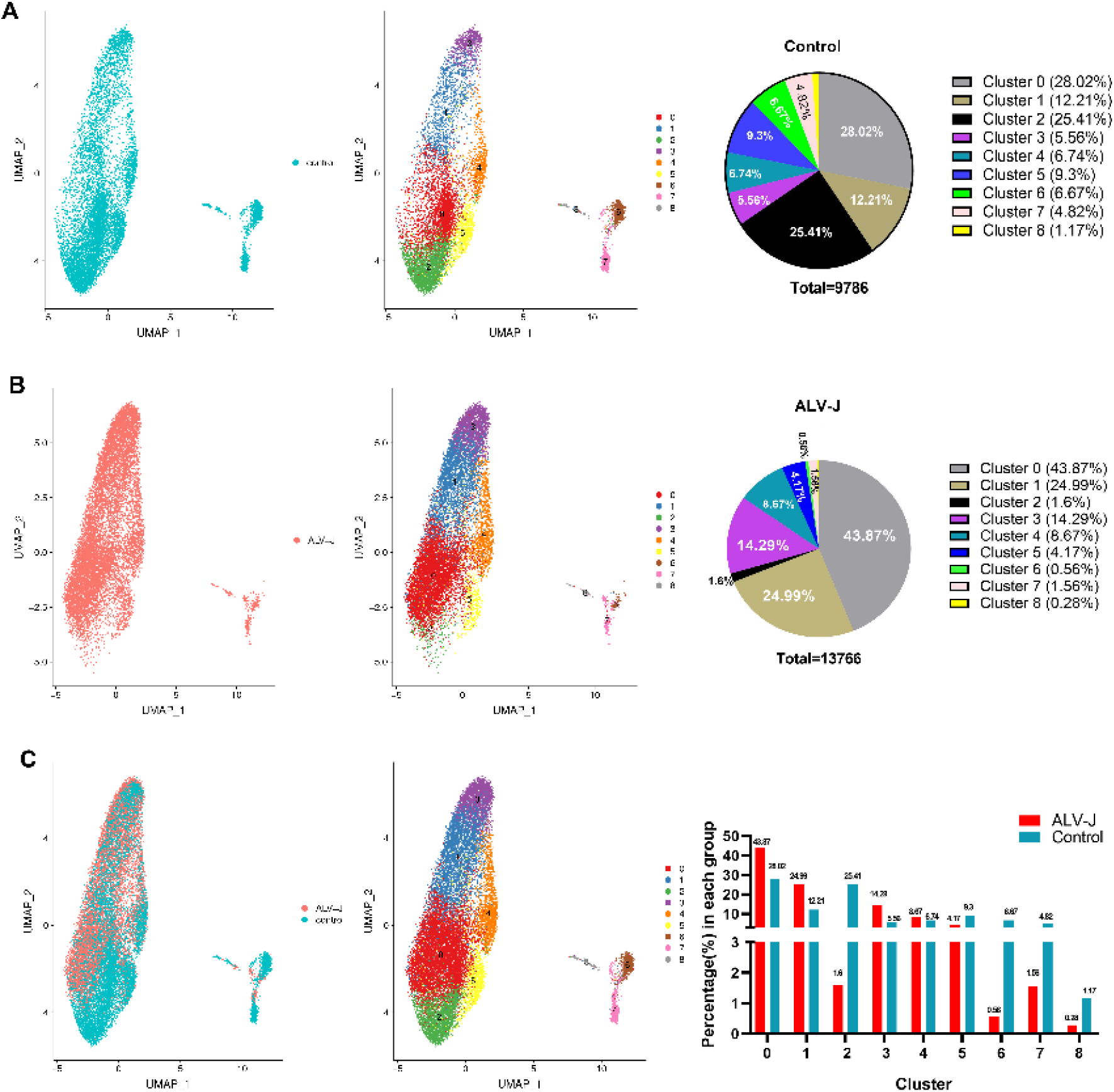
Single-cell profiling of cell populations in chicken peripheral blood lymphocytes (PBLs) collected from avian leukosis virus subgroup-J (ALV-J)-infected and control chickens at 21 days post infection (DPI). **(A)** Uniform Manifold Approximation and Projection (UMAP) displaying all cell clusters and their percentage in the control PBLs. **(B)** UMAP displaying all cell clusters and their percentage in PBLs from ALV-J infected chicken. **(C)** Cell distribution and percentage of each cluster in PBLs from ALV-J infected and control chickens.

For further analyzing the immune signatures of each cluster in chicken PBL, we identified the up-regulated DEGs in each cluster and analyzed the DEGs enriched in the GO terms “immune system process”, “response to stimulus”, and “defense response to virus”. The results showed that immune-related DEGs were largely detected in clusters 6, 7, and 8 (Fig. 2B and files S2-S3), which implied that, in PBL, these clusters are the main effector responses to pathogenic stimuli. Additionally, the expression levels and the percentage of cells expressing the top five genes in each cluster are shown in a dot plot (Fig. 2C, file S4, and Fig. S1); these need to be confirmed in future research and are proposed to be used as marker genes for each cluster of chicken PBLs.

**Figure 2.**
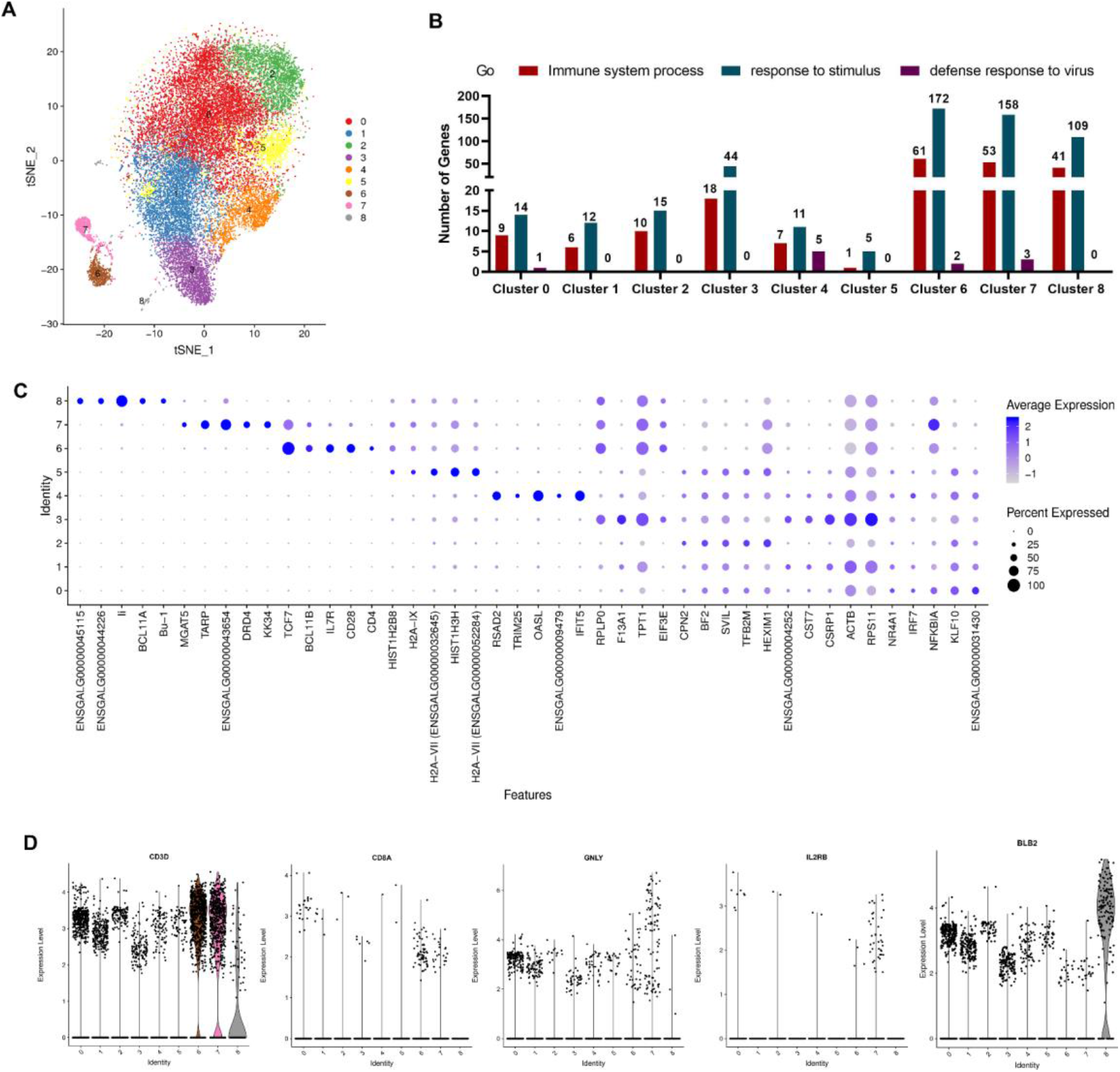
Analysis of cell types of each cluster in chicken PBLs. **(A)** t-Distributed Stochastic Neighbor Embedding (t-SNE) projection representing the eight clusters of cells identified in the chicken PBL pools (unified set of control and ALV-J infection samples). **(B)** The statistics of genes involved in the GO terms “immune system process”, “response to stimulus” and “defense response to virus”, as analyzed in each cluster. **(C)** Top five DEGs (x-axis) identified in each cluster (y-axis). Dot size represents the fraction of cells in the cluster that express the gene; intensity indicates the mean expression (Z-score) in the expressing cells, relative to other clusters. (D) Expression levels of characteristic marker genes (CD3D, CD8A, GNLY, IL2RB, and BLB2) in PBL clusters.

Based on the expression of classical CD3 marker, we could define cluster 6 and cluster 7 as T cells (Fig. 2D). Cluster 6 mainly included activated CD4^+^ T cells (CD3^+^CD4^+^IL7R^+^CD28^+^). KK34 is reported to encode an IL-5-like transcript that was specifically expressed by avian γδT cells, which may mediate T helper 2 (Th2)-cytokine-dependent allergy [27]. Meanwhile, other studies reported that dopamine receptor (DRD4) is involved in Th2 cell differentiation and inflammation [28]. Therefore, we defined cluster 7 as Th2-like γδT cell (CD3^+^KK34^+^DRD4^+^; Fig. 2C and D). In addition, we found that a few cytotoxic CD8^+^ T cells (CD3^+^CD8^+^GNLY^+^) mixed in clusters 6 and 7 (Fig. 2D). Therefore, we planned to regroup clusters 6 and 7 for a more detailed display of data in the following analysis. Additionally, cluster 8 should be B cells according to the expression of the known gene markers, BCL11A and Bu-1 (ENSGALG00000015461) (Fig. 2C). Interestingly, the top five genes in cluster 4 were mainly interferon stimulating genes (ISGs), including RSAD2, TRIM25, OASL, and IFIT5, which implied that cluster 4 also contained a type of important antiviral immune cell (Fig. 2C). Therefore, we defined cluster 4 as ISG expressing cells in PBL, which needs to be further verified in future studies. Unfortunately, we were unable to define clusters 0, 1, 2, 3, and 5 based on the few known chicken cell-type markers and the top five genes expressed by cells in these clusters.

### Most DEGs were detected in the T cell population (cluster 6 and cluster 7) in response to ALV-J infection at 21 DPI

We calculated the DEGs between cell populations of PBLs from the ALV-J infected and control chickens at 21 DPI using Seurat. It was found that the total number of DEGs and the DEGs enriched in the GO terms “response to stimulus” and “cell proliferation” were largely detected in clusters 6 and 7 (Fig. 3A and file S5), indicating that cells in clusters 6 and 7 might have played an important role during the antiviral response to ALV-J in the infected chicken. We further discovered that the decreased percentage of clusters 6 and 7 in PBLs from ALV-J infected chicken might be a result of the up-regulated pro-apoptotic factors, ITPR2 [29] and JARID2 [30] (Fig. 1C, Fig. 3B, and file S5). Next, an immune related DEG analysis was performed after regrouping clusters 6 and 7 in the study outlined below.

**Figure 3.**
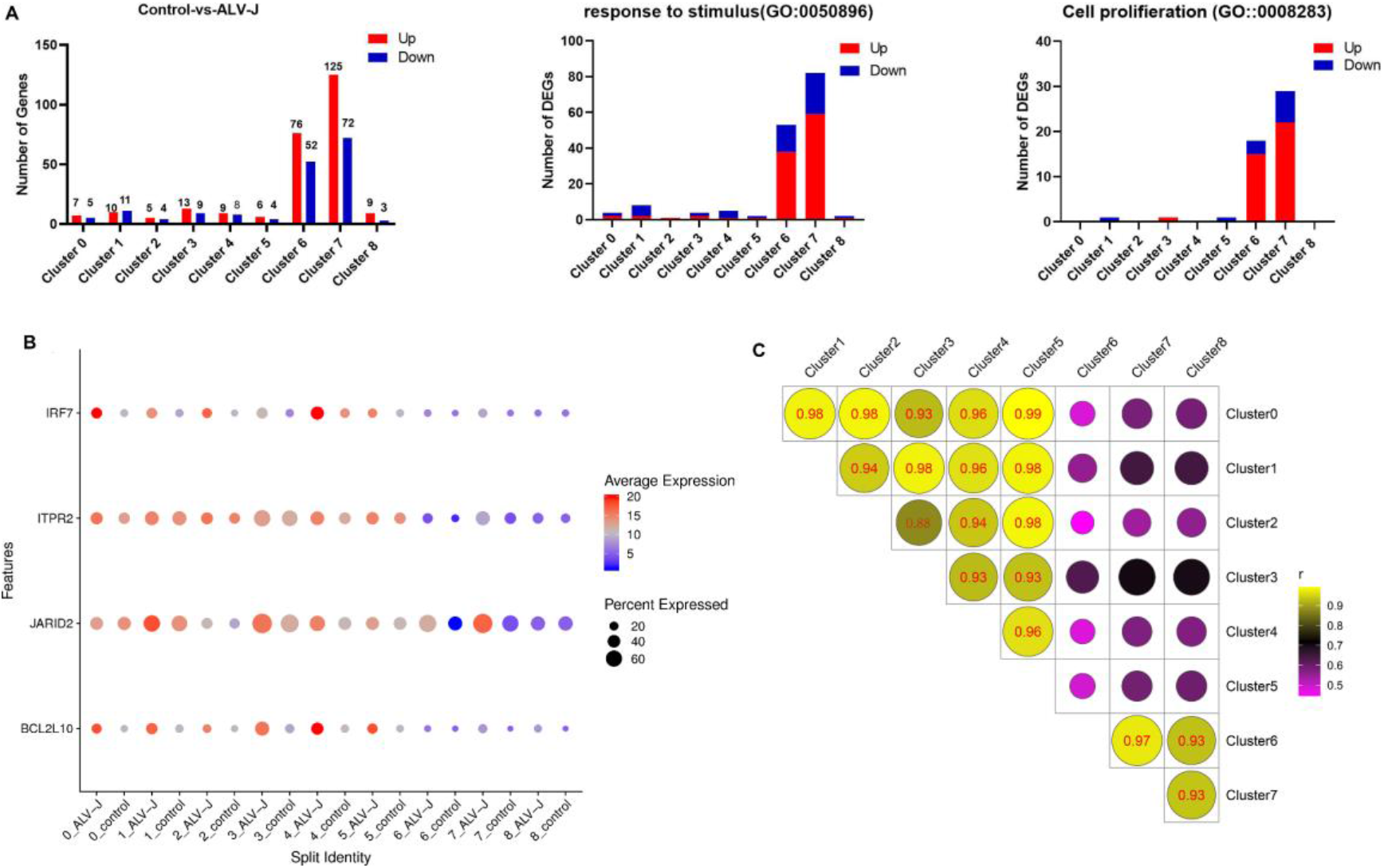
Differentially expressed genes (DEGs) analysis in PBLs from ALV-J infected and control chickens within cell clusters. **(A)** Histogram showing all up-regulated (red) and down-regulated (green) DEGs, and the number of DEGs involved in the GO terms “response to stimulus” and “cell proliferation” in PBLs from ALV-J infected chicken compared to control PBLs within each cluster. **(B)** Dot plot representing selected DEGs (IRF7, ITPR2, JARID2, and BCL2L10) expressed in eight clusters within which cells from the ALV-J infected sample were compared with the control sample. The intensity represents the expression level, while dot size represents the percentage of cells expressing each gene. **(C)** Pearson’s Correlation analysis of different cell clusters based on gene expression levels.

Meanwhile, some DEGs in other clusters were analyzed. We found that an important immune gene, interferon regulator 7 (IRF7), revealed up-regulation in clusters 0, 1, 2, and 3 in response to ALV-J infection at 21 DPI (Fig. 3B and file S5). We also found that the anti-apoptotic gene BCL2L10 [31] was up-regulated in clusters 0, 1, 3, 4, and 5 of PBLs from ALV-J infected chicken (Fig. 3B and file S5). Besides, Pearson’s correlation analysis indicated that clusters 0, 1, 2, 3, 4, and 5 were strongly correlated between each other, but displayed very low correlation with clusters 6, 7, and 8 (corresponding to T cells and B cells; Fig. 3C), which implied that the function of cells in clusters 0, 1, 2, 3, 4, and 5 may be related to innate immune responses during early infection stage.

### T cell populations of clusters 6 and 7 could be further divided into four distinct cell clusters in the PBLs

The above-mentioned T cell populations of clusters 6 and 7, both in PBLs from the ALV-J infected or control chicken, were further divided into four distinct clusters (named as clusters A0 to A3) and visualized using UMAP (Fig. 4A-C). Compared to those in the control PBLs, the percentages of clusters A0 and A1 in the PBLs from ALV-J infected chicken was decreased. Conversely, the percentage of clusters A2 and A3 was significantly increased in PBLs from ALV-J infected chicken (Fig. 4C). Next, the top five genes expressed in clusters A0 to A3 were picked as potential marker genes and they are shown in a dot plot (Fig. 4D, Fig. S2, and File S6). Judging according to the classical marker genes and our above analysis, we considered cluster A0 as activated CD4^+^ T cells (CD3^+^CD4^+^IL7R^+^CD28^+^), cluster A3 as cytotoxic CD8^+^ T cells (CD3^+^CD8^+^IL2RB^+^GNLY^+^), and cluster A1 as Th2 like γδT cells (CD3^+^KK34^+^DRD4^+^) (Fig. 4D and E).

**Figure 4.**
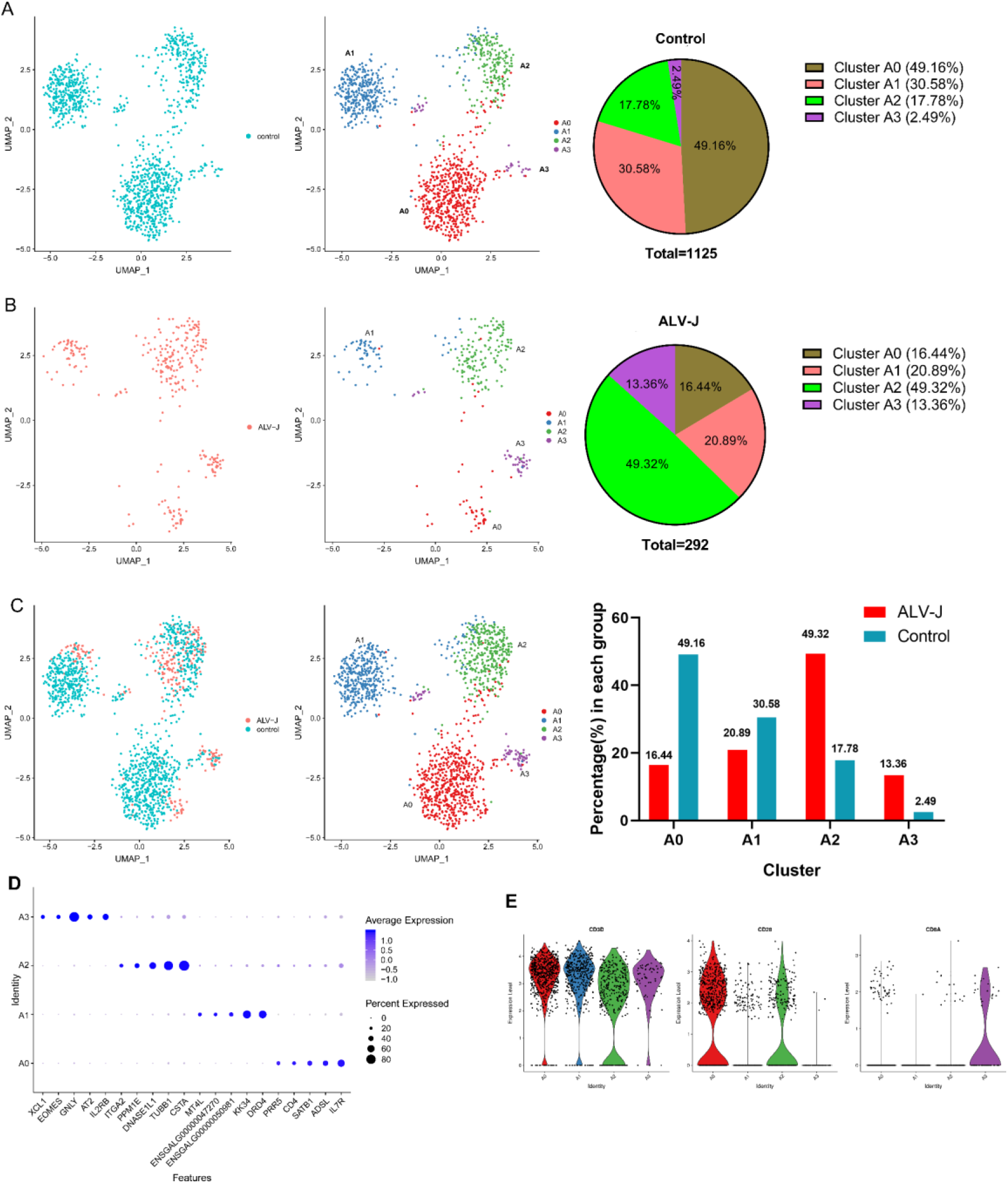
T cell population of clusters 6 and 7 were divided into four distinct cell clusters in the PBLs. **(A)** UMAP displaying the cell clusters and their percentage of the regrouped T cell populations (clusters 6 and 7) in the control PBLs. **(B)** UMAP displaying the cell clusters and their percentage of the regrouped T cell population (clusters 6 and 7) in PBLs from ALV-J infected chicken. **(C)** The regrouped T cell distribution and percentage of each cluster in PBLs from ALV-J infected and control chickens. **(D)** Top 5 DEGs (x-axis) identified in clusters A0–A3 (y-axis). **(E)** Expression levels of characteristic marker genes (CD3D, CD28, and CD8A) in clusters A0–A3.

Although we were unable to define cluster A2 based on their top five genes and a few known markers of various chicken cell-types, using pseudotime analysis, we believe that clusters A0 (activated CD4^+^ T cells, namely Th0 cells), A1 (Th2-like cells), and A2, in both ALV-J infected and control samples, demonstrated a potential differentiation correlation (Fig. 5A). Cluster A0 (Th0 cells) is mainly located at the early stage of the pseudo-time trajectories, whereas clusters A2 and A1 (Th2-like cells) are mainly located at the late stage (Fig. 5A and B). These results suggest that clusters A2 and A1 are probably differentiated from A0 and imply that cluster A2 may represent Th1-like cells. Interestingly, cells in the control PBL are mainly distributed in the A0 (Th0) and A1 (Th2-like) cell clusters (Fig. 5C-E). Conversely, ALV-J-infected PBL are highly enriched in the terminally differentiated A2 cell clusters (Th1-like; Fig. 5C-E), which indicates that ALV-J infection induced CD4^+^ T cell activation and differentiation into the Th1 phenotype. Finally, the branch-dependent differential gene hierarchy clustering heat map is shown in supplementary Fig. 3; it contains the top 10 branching DEGs displayed in Fig. 5F. Of note, BRT-1, CSTA, ENSGALG00000046729, HPSE, IFI6, ITGB3, and TUBB1 may be associated with Th1 cell differentiation (Fig. 5D-F).

**Figure 5.**
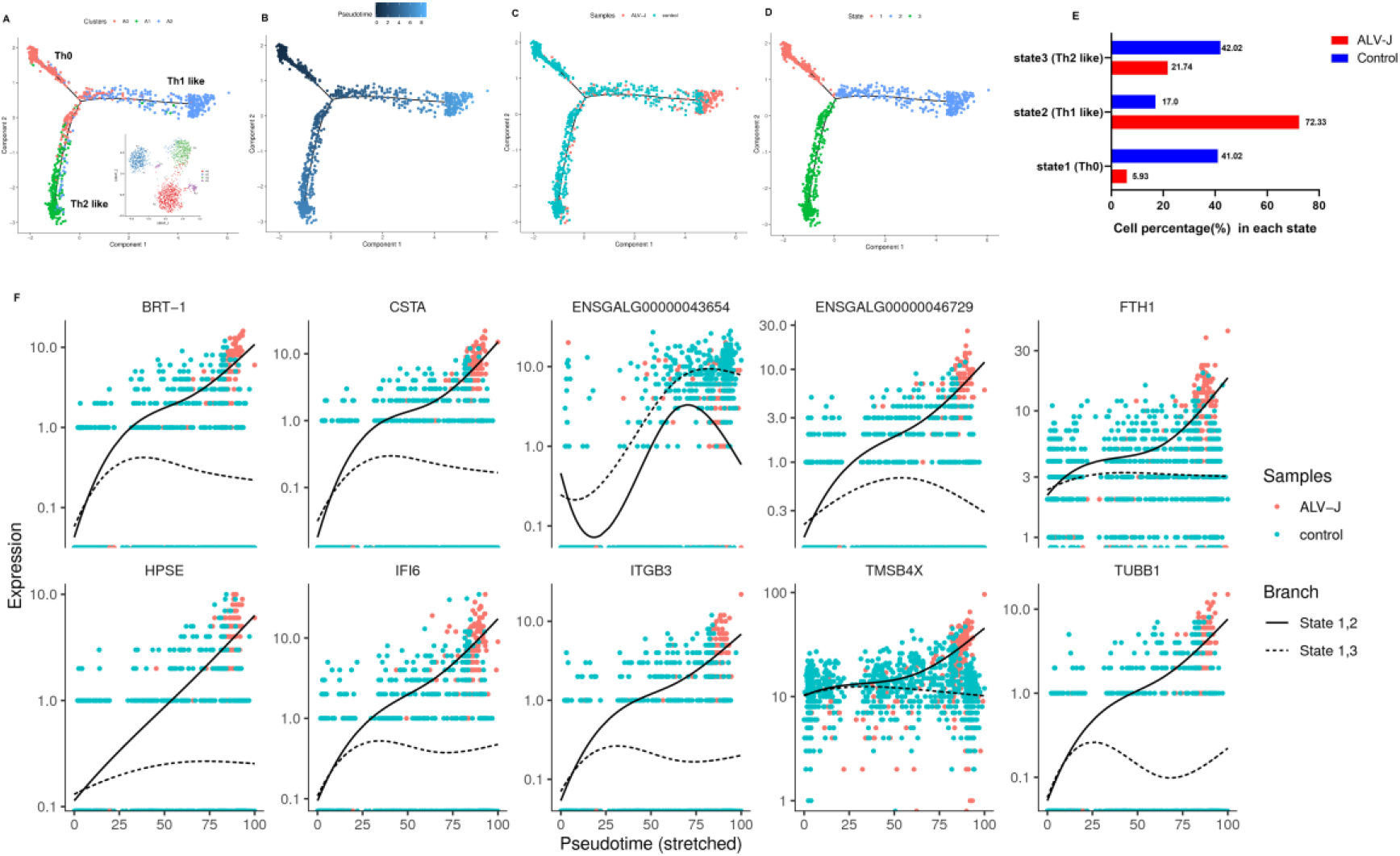
Pseudotime analysis of clusters A0, A1, and A2. **(A)** Mapping of clusters A0 (Th0), A1 (Th2-like) and A2 (Thl-like) to the pseudotime trajectory. **(B)** Pseudotime trajectory calculated from all cells of clusters A0–A2 in the control and ALV-J infected samples. Darker colored dots represent a shorter pseudotime and earlier differentiation period. **(C)** Mapping of cells in control and ALV-J infected samples to the pseudotime trajectory. **(D)** The cell states of pseudotime trajectory partitioned from all cells of clusters A0–A2 in the control and ALV-J infected samples. **(E)** Cell percentage of each state between the ALV-J infected and control samples. **(F)** Dynamics of the top 10 branching DEGs. Full line: state 1, 2; imaginary line: state 1, 3.

### Cluster A2 (Th1-like) is the vital cell type in response to ALV-J infection

Our previous results show that clusters 6 and 7 play a potentially important role in the antiviral response during ALV-J infection based on their largely immune-related DEG expression (Fig. 3A and file S5). Meanwhile, the T cell populations in clusters 6 and 7 could be grouped into four distinct sub clusters (Fig. 4). To further investigate each T cell population response, we firstly calculated the DEGs between corresponding T cell population of the PBLs from ALV-J infected and control chickens at 21 DPI using Seurat. Strikingly, the highest numbers of total DEGs and the DEGs enriched in the GO terms “response to stimulus” and “cell proliferation” were predominant in cluster A2 (Th1-like T cells; Fig. 6A and file S7), suggesting that cluster A2 represents likely the vital effectors among PBLs of the ALV-J infected host.

**Figure 6.**
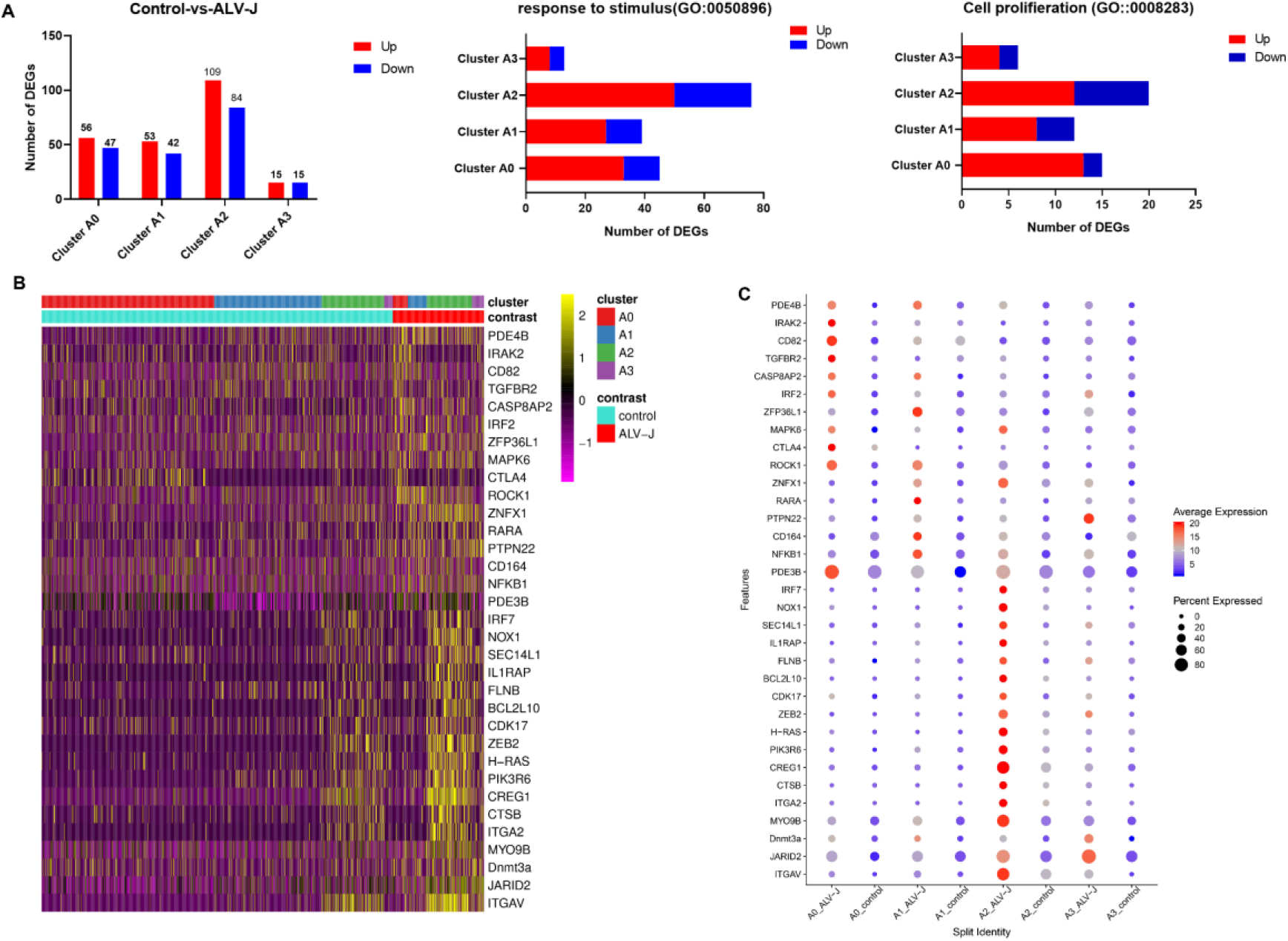
Landscape of immune-related gene expression in clusters A0–A3 between the ALV-J infected and control samples. **(A)** Histogram showing all the up-regulated (red) and down-regulated (green) DEGs, and the number of DEGs involved in GO terms “response to stimulus” and “cell proliferation” in ALV-J stimulated cells compared to control cells in clusters A0–A3. **(B)** Heatmap showing the normalized expression (Z-score) of all immune-related DEGs in various cells of clusters A0–A3 within which cells from the ALV-J infected sample are compared with the control sample. **(C)** Dot plot representing DEGs expressing in clusters A0–A3 within which cells from the ALV-J infected sample is compared with the control sample. The intensity represents the expression level, while the size of the dots represents the percentage of cells expressing each gene.

Next, the specific DEGs of the four sub clusters enrichment in “ response to stimulus” (GO:0050896) were exhibited using a volcano plot (Fig. S4). Moreover, a total of 33 important immune-related genes were screened from clusters A0 to A3 and their expression levels were presented as a heat map and a dot plot (Fig. 6B-C and file S7). We also observed that most of these immune-related genes were highly expressed in cluster A2. It is worth noting that BCL2L10, H-RAS, IRF7, NOX1, SEC14L1, IL1RAP, FLNB, CDK17, ZEB2, PIK3R6, CREG1, CTSB, and ITGA2 were up-regulated in cluster A2, rather than in the other three clusters (Fig. 6C and file S7). Moreover it is reported that H-RAS act as critical controllers of Th1 responses via transmitting TCR signals for the Th1 priming of CD4^+^ T cells [32], which further supports our previously described results that cluster A2 may represent Th1 cells. Besides, we found that PDE3B, RARA, and CD164 were merely up-regulated in cluster A1, while IRAK2, CD82, IRF2, MAPK6, TGFBR2, and CTLA4 were only up-regulated in cluster A0. Furthermore, the apoptosis-associated gene, CASP8AP2 [33], exhibited increased expression in clusters A0 and A1, which could potentially explain their proportional decrease in PBL after infection. Taken together, these up-regulated genes could potentially serve as marker genes for chicken Th1-like cells (cluster A2), Th2-like cells (cluster A1) and Th0 cells (cluster A0), respectively.

Finally, we conducted an interaction network analysis of the 33 candidate DEGs based on the STRING database (Fig. 7). The results implied that the 13 DEGs marked as red were likely the more important hub genes. Specifically, 4 hub genes including ITGA2, IL1RAP, NOX1 and CDK17 were up-regulated in cluster A2 (Th1-like cells, Fig. 6C and file S7). 3 hub genes including IRAK2, CTLA4 and TGFBR2 were up-regulated in cluster A0 (Th0 cells, Fig. 6C and file S7). 2 hub genes including MYO9B and ZNFX1 were up-regulated in Th1-like cells and cluster A1 (Th2-like cells, Fig. 6C and file S7). Besides, identified hub gene PDE3B was up-regulated in Th2 like cell; ITGAV was up-regulated in Th1-like, Th2-like and cytotoxic CD8^+^ T cell populations; CASP8AP2 was up-regulated in Th0 and Th2-like cells; JARID2 was up-regulated in Th0, Th1-like and cytotoxic CD8^+^ T cell populations (Fig. 6C and file S7).

**Figure 7.**
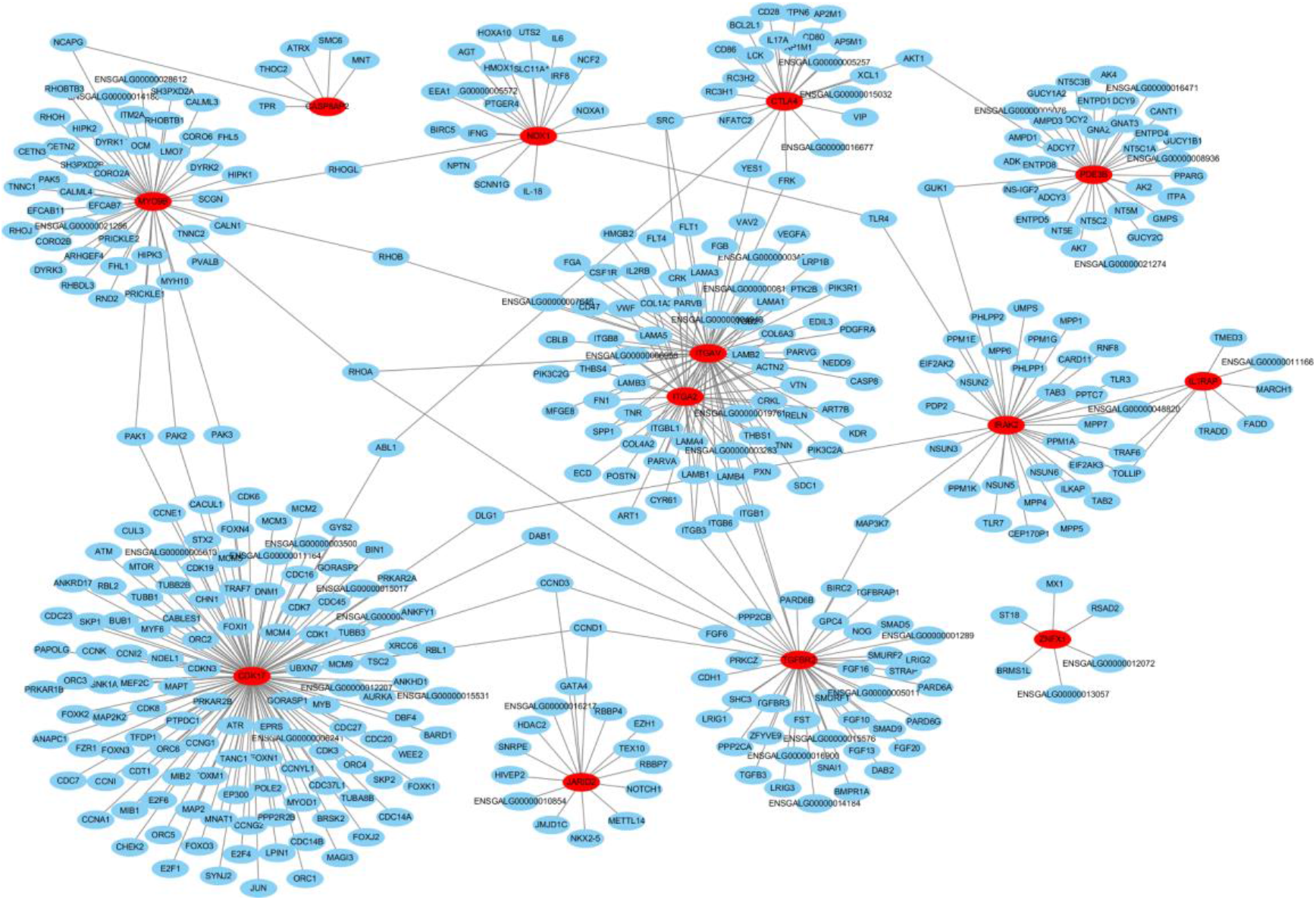
The interaction network analysis of the candidate 33 DEGs based on the STRING database. In this network, red nodes represent hub genes, and lines represent the associations.

## Discussions

Adaptive immunity, including T cell and B cell response, is the foundation upon which vaccines are developed. Despite decades of research, we still have limited insights into the ability of the avian immune response to eliminate pathogens at the cellular and molecular levels. For example, antibody level and T cell proportional change in the peripheral blood are usually used to evaluate a virus-induce immune response [3,5,13,34]. Flowcytometry-based phenotyping and functional evaluation of chicken T cells are limited by reagents and methods availability [1]. Fortunately, advances in scRNA-seq can assist with these problems and significantly promote our understanding of the molecular details regarding efficient immune responses to viral infections. Previously, we found that ALV‐J viremia was eliminated by 21 DPI when an up-regulated CD8^+^ T cell ratio and low antibody levels were detected [3]. In the current study, we used 10x scRNAseq on PBLs in ALV-J infected chicken and uninfected chicken (control) at 21 DPI to characterize two major aspects of immune response against the invading pathogen: the immune cell composition and their responses to infection.

In this work, eight distinct cell clusters in chicken PBLs were identified following analysis of scRNA-seq data. Strikingly, we found that T cell populations (clusters 6, 7) and B cell populations (cluster 8) occupied a very small percentage in chicken PBLs. Of note, ALV-J infection has an obvious impact on the cell composition of PBLs. Specifically, the total number and percentage of T cells, B cells, and cluster 2 were distinctly reduced in PBLs from ALV-J infected chicken compared to the control PBLs. Furthermore, up-regulated ITPR2 [29] and JARID2 [30] expression may be involved in T cell apoptosis in PBLs from ALV-J infected chicken (Fig. 1C, Fig. 3B and file S5). On the other hand, the percentage of clusters 0, 1, and 3 were obviously increased in PBLs from ALV-J infected chicken compared to the control PBLs, which may be associated with the up-regulation of the anti-apoptotic gene, BCL2L10 [31]. Additionally, Pearson’s correlation analysis showed that clusters 0, 1, 2, 3, 4, and 5 exhibited strong correlation, but they displayed very low correlation with the T cell and B cell populations (clusters 6, 7, and 8; Fig. 3 C). Unfortunately, clusters 0, 1, 2, 3, 4, and 5 were unable to be defined based on the few known chicken cell-type markers and five most highly expressed genes in these clusters. Interestingly, the top expressed genes in cluster 4 were mainly chicken ISGs including RSAD2, TRIM25, OASL, and IFIT5 [35], which implied that cluster 4 contained important antiviral immune cells (Fig. 2C). Moreover, IRF7 which is involved in IFN-β signaling and ISGs inducer [35,36], revealed up-regulation in clusters 0, 1, 2, and 3 response to ALV-J infection at 21 DPI (Fig. 3B, file S5). According to these results, the function of clusters 0, 1, 2, 3, 4, and 5 may be related to innate immune response at the early infection stage. The specific cell types and function of these clusters need to be further confirmed in future studies.

Most DEGs were detected in the T cell population (cluster 6 and cluster 7) response to ALV-J infection at 21 DPI (Fig. 3A and file S5). Furthermore, the T cell population (cluster 6 and cluster 7) could be grouped into four distinct cell clusters including activated CD4^+^ T cells (Cluster A0), Th2-like γδT cell (Cluster A1), Th1-like cell (Cluster A2), and cytotoxic CD8^+^ T cells (Cluster A3), which were defined based on the expression of marker genes and pseudotime analysis results. Here, pseudotime analysis implied that activated CD4^+^ T cells in chicken PBLs could become terminally polarized to a Th1 or Th2 phenotype. Moreover, ALV-J infection induced CD4^+^ T cell activation and differentiation into a Th1 phenotype was probably associated with expression of the top 10 branching DEGs (Fig. 5D-F). In addition, the Th1-like cell population (Cluster A2) was vital in response to ALV-J infection at 21 DPI based on the expression of largely immune-related DEGs. Compared to the control PBLs, the percentage of Th1-like cells (cluster A2) and cytotoxic CD8^+^ T cells (cluster A3) were increased in the T cell population of PBLs from ALV-J infected chicken. It is also known that Th1 cells can help cytotoxic CD8^+^ T cell survival, activation, and memory response [37]. Conversely, B cell percentage was decreased, and few DEGs were detected after ALV-J infection (Fig. 1C and Fig. 3A). It is also reported that ALV-J infection inhibits the proliferation, maturity, and response of B cells [38]. Therefore, it was the T cell response, including Th1-like cells and cytotoxic CD8^+^ T cells, instead of the B cell response that eliminated ALV‐J viremia at 21 DPI. Our previous animal experiments also verified that T cell response as opposed to humoral immunity was the key factor defending against ALV-J infection [3].

More importantly, we identified 13 hub genes for the first time that were up-regulated in each chicken T cell population after ALV-J infection. Of note, CDK17, reported to inhibit porcine reproductive and respiratory syndrome virus (PRRSV) infection [39]; Nox1, reported to suppress influenza A virus induced lung inflammation and oxidative stress [40]; and IL1RAP, reported to give negative regulation of Transmissible gastroenteritis virus (TGEV) induced mitochondrial damage [41], were all up-regulated in Th1-like cells of PBLs from ALV-J infected chicken. The information reminded us that IL1RAP, NOX1, and CDK17 can be potential functional marker genes of Th1-like cells exerting antiviral function. Furthermore, we found that ZNFX1 was up-regulated in Th1-like and Th2-like cells, and was involved in inducing IFN and ISG expression [42], which indicated the complexity of antiviral immunity in T cells. On the other hand, IRAK2, reported to potentially suppress avian infectious bronchitis virus (IBV) infection [43], exhibited up-regulated expression in activated CD4^+^ T cells (cluster A0, Th0 cell) of PBLs from ALV-J infected chicken. But, interestingly, CTLA4, a critical co-receptor for Treg cell function [44,45], and TGFBR2, the important TGF-β receptor suppressing proliferation and terminal differentiation of antiviral CD4^+^ T cells [46], also exhibited up-regulated expression in activated CD4^+^ T cells. In the current study, we did not identify the two subpopulations of chicken regulatory T cells (Treg cells) including TGF-beta^+^CD4^+^ T cells and CD4^+^CD25^+^ T cells [47,48]. We hypothesize that Th0 cells may negatively regulate T cell response to homeostatic control through CTLA4 and TGFBR2. In addition, further studies are needed to confirm whether or not Th0 cells are able to differentiate into regulatory T cells. Therefore, further scRNA-seq experiments with additional pathogens and at a greater number of time-points are needed in order to fully uncover the global and pathogen specific cell-type immune responses.

In summary, our scRNA-seq study based on PBLs in chickens produced a rich data resource that can be mined during future experiments to address the function of these cells throughout development and in response to pathogenic infection. The “marker genes” that were assigned to different clusters need to be verified with specific antibodies. To the best of our knowledge, the hallmark genes for each T cell population involved in ALV-J infection were identified here for the first time. Moreover, using pseudotime analysis, we found that chicken CD4^+^ T cells could differentiate into Th1-like and Th2-like cells. With respect to the control PBLs, ALV-J infection had an obvious impact on PBL cell composition. B cells were decreased and inconspicuous in response in PBLs from ALV-J infected chicken at 21 DPI. Cytotoxic Th1-like cells and CD8^+^ T cells are potential key factors in the defense against ALV-J infection.

## Acknowledgments

Grateful thanks are given to the South China Agricultural University high-level talent launch project and “Fuji Peiyou” project of College of Veterinary Medicine, South China Agricultural University. We are extremely appreciative of the help given by Gene Denovo Corp. during bioinformatics analysis. Special thanks are given to Jingjing Ning for help in project coordination.

## Funding

This work was supported by the National Natural Science Foundation of China (31802174 and 31830097) and China Postdoctoral Science Foundation Grant (2019T120735), 111 Project (D20008).

## Author contributions

MMD and MF participated in the design of the study, performed the experiments, collected and analyzed data, and drafted the manuscript. ZWL assisted with data analysis. WSC and ML participated in the coordination of the study and revised the manuscript. All authors read and approved the final manuscript.

## Competing financial interests

The authors declare that they have no conflicts of financial interest.

## Data availability

Sequencing data have been deposited in BioProject under the accession number PRJNA687808.

---

**Supplementary Fig.1 (Fig.S1). t-SNE displaying top 5 DEGs identified in clusters 0–8 of PBLs**

**Supplementary Fig.2 (Fig.S2). t-SNE displaying top 5 DEGs identified in clusters A0–A3 of PBLs**

**Supplementary Fig.3 (Fig.S3). Heat map of branch-dependent differential gene hierarchy clustering**

**Supplementary Fig.4 (Fig.S4). Volcano plot of identified DEGs involved in the GO term “response to stimulus” (GO:0050896) between ALV-J infected and control samples in clusters A0–A3**

**Supplementary file 1 (file S1). scRNA-seq data statistics.**

**Supplementary file 2 (file S2). Up-regulated differentially expressed genes (DEGs) in each cluster of PBLs.**

**Supplementary file 3 (file S3). Up-regulated DEGs involved in “immune system process”, “response to stimulus” and “defense response to virus” GO terms in each PBL cluster.**

**Supplementary file 4 (file S4). Top 5 DEGs of each chicken PBL cluster.**

**Supplementary file 5 (file S5). DEGs found in each cluster between the ALV-J infected and control samples.**

**Supplementary file 6 (file S6). Top 5 DEGs in clusters A0–A3**

**Supplementary file 7 (file S7). DEGs in clusters A0–A3 between the ALV-J infected and control samples.**

## Notes

### Competing Interest Statement

The authors have declared no competing interest.

